# MARCH5-dependent NLRP3 ubiquitination is an essential step for NEK7 docking on the mitochondria

**DOI:** 10.1101/2023.01.12.523764

**Authors:** Yeon-Ji Park, Niranjan Dodantenna, Tae-Hwan Kim, Ho-Soo Lee, Young-Suk Yoo, Eun-Seo Lee, Jae-Ho Lee, Myung-Hee Kwon, Ho Chul Kang, Jong-Soo Lee, Hyeseong Cho

## Abstract

The NLRP3 inflammasome is a global immune-sensor that is activated by a repertoire of endogenous and exogenous stimuli. NLRP3 translocates to mitochondria but whether mitochondria involvement in the inflammasome assembly is unclear. Here, we show that the mitochondrial E3 ligase MARCH5 is a key regulator of NLRP3 inflammasome assembly. Myeloid cell-specific *March*5 conditional knockout (*March5* cKO) mice exhibited an attenuated mortality rate upon LPS or *Pseudomonas aeruginosa* challenge. Macrophages derived from *March5* cKO mice failed to secrete IL-1β and IL-18 after microbial infection. Mechanistically, MARCH5 interacts with the NACHT domain of NLRP3 and promotes K27-linked polyubiquitination of K324 and K430 residues of NLRP3. Ubiquitination-defective NLRP3 mutants neither bind to NEK7, nor form NLRP3 oligomers, but remain binding to MAVS. Accordingly, NLRP3 mutants led to abortive ASC speck formation and diminished IL-1β production. We propose that MARCH5-dependent NLRP3 ubiquitination creates a docking site for NEK7 binding, playing as a fundamental step-wise regulator on the mitochondria.

## Introduction

Activation of innate immune system is initiated by the recognition of molecules derived from pathogens (PAMPs: pathogen-associated molecular patterns) (Takeuchi & Akira, 2010) or the damaged host cells (DAMPs: damage-associated molecular patterns) (Chen & Nunez, 2010). NLRP3 (NOD-like receptor family pyrin domain containing 3) is one of the principal pattern recognition receptors (PRRs) that recognize diverse array of PAMPs and DAMPs (Ratsimandresy *et al*, 2013). After the recognition, NLRP3 may undergo conformational change and interact with an adapter protein, ASC (apoptosis-associated speck-like protein) and the effector molecule pro-caspase-1 to form the large protein complex of the NLRP3 inflammasome (Guo *et al*, 2015). Apparently, two sequential steps are needed for the formation of active NLRP3 inflammasome: priming and activation. Priming involves transcriptional upregulation of TNF-*α*, IL-6 and NLRP3 genes as well as the inactive cytokine precursor genes pro-IL-1β and pro-IL-18 through the activation of upstream receptors such as TLR4 (Toll-like receptor 4). The NLRP3 inflammasome assembly and its activation are triggered by chemically and structurally diverse stimuli, including potassium efflux, ATP, nigericin, monosodium urate (MSU), and others, causing the activation processes to be complex and less obvious. Complete inflammasome assembly triggers the self-cleavage of pro-caspase-1 and the formation of active caspase-1, converting pro-IL-1β and pro-IL-18 to their mature forms for release (Guo *et al*., 2015). Notably, none of the stimuli bind to NLRP3 directly (Kelley *et al*, 2019). In fact, dysregulation or persistent NLRP3 signaling underlies various inflammatory diseases and autoimmune disorders (Fusco *et al*, 2020). Uncovering the underlying mechanism would enable the development of targeted therapeutic approaches to control these diseases.

Mitochondria are intensely involved in NLRP3 inflammasome activation. NLRP3 stimuli increase the production of ROS (reactive oxygen species) and release oxidized fragments of mitochondrial DNA into the cytosol, both of which stimulate inflammasome activation (Cruz *et al*, 2007; Heid *et al*, 2013; Shimada *et al*, 2012b). In addition, LPS-induced mitochondrial DNA synthesis has also been shown to contribute to NLRP3 inflammasome activation (Zhong *et al*, 2018). Increasing evidences have shown how damaged mitochondrial DNA and RNAs are released into the cytosol and stimulate NLRP3 and other inflammatory signaling (Shimada *et al*, 2012a; Tigano *et al*, 2021). The NLRP3 localization is not limited to the cytosol but is also observed in mitochondria, ER and Golgi complex. Meanwhile, the mitochondria-associated adaptor molecule MAVS mediates the recruitment of NLRP3 to the mitochondria, which is necessary for the activation of the NLRP3 inflammasome (Subramanian *et al*, 2013). Similarly, mouse NLRP3 and caspase-1 independently interact with the mitochondrial lipid cardiolipin (Iyer *et al*, 2013). However, it is still unclear why mitochondrial translocation of NLRP3 is required for NLRP3 inflammasome activation. Another important step for the NLRP3 activation is NLRP3 oligomerization. In the resting state, inactive NLRP3 retains its folded structure, but it converts to the unfolded form and undergoes self-oligomerization upon activation. Accumulating evidences suggest that NLRP3 oligomerization requires its binding to NEK7 (NIMA-related kinase 7), which forms a bridge between adjacent NLRP3 subunits (Sharif *et al*, 2019). It is still unknown how NLRP3-NEK7 in cells interaction is operated.

MARCH5 (also known as MITOL) is an E3 ubiquitin ligase that localizes to the mitochondrial outer membrane (Bauer *et al*, 2017). MARCH5 plays a central role in maintaining mitochondrial and cellular homeostasis by regulating proteins responsible for mitochondrial dynamics and protein aggregation accumulation in the mitochondria (Nagashima *et al*, 2014; Yonashiro *et al*, 2006). MARCH5-mediated ubiquitination of target proteins is crucial for ER-mitochondria contact and protein import into the mitochondria (Phu *et al*, 2020; Sugiura *et al*, 2013; Takeda *et al*, 2019). Lack of MARCH5 function in mitochondrial dynamics induces cellular senescence or affects cell survival (Park *et al*, 2010; Shiiba *et al*, 2021). Its absence also aggravates neuronal pathogenesis (Takeda *et al*, 2021). In addition, MARCH5 serves as an important immune regulator and a positive regulator of TLR7 signaling by transferring K63-linked polyubiquitin chains to TRAF family member-associated NF-κB activator (TANK) (Shi *et al*, 2011). MARCH5 also targets massive complexes of activated RIG-I-MAVS oligomers and dissolves them to prevent persistent immune activation (Park *et al*, 2020; Yoo *et al*, 2015).

This study shows that MARCH5 is a cue for NLRP3 oligomerization on mitochondria. MARCH5 interacts NLRP3-MAVS complex and transfers K27-linked polyubiquitin to NLRP3, which provides a docking site for NEK7 binding to NLRP3, initiating NEK7-NLRP3 multimeric complex formation. Our findings highlight the mitochondria as an onset platform for NLRP3 immune complex formation.

## Results

### Myeloid cell-specific *March5* conditional knockout (*March5* cKO) mice exhibit an attenuated mortality rate and inflammatory response to septic shock

Mitochondria are involved in the cellular innate immune response against viruses. The mitochondrial outer membrane provides a surface platform for RLR signaling against the cytosolic viral RNA genome (Banoth & Cassel, 2018). Our previous studies revealed that MARCH5 degrades the active form of prion-like MAVS aggregates and oligomerizes RIG-I, preventing persistent immune reactions (Park *et al*., 2020). Here, we addressed whether MARCH5 has any immunoregulatory effects upon bacterial infection. We generated myeloid cell-specific *March5* knockout C57BL/6 mice to address this issue using the Cre-loxP recombination system. To selectively delete the *March5* gene in myeloid cells, we crossbred *March5*^*fl/fl*^ mice with Lys2-Cre mice, deleting exon 3 of the *March5* gene with Cre recombinase (Fig EV1A). We examined the mRNA levels using primers targeting exon 3 of the *March5* gene in several tissues, namely, the brain, heart, spleen, and tail as well as in bone marrow-derived macrophages (BMDMs) derived from *March5*^*fl*/*fl*^ and *March5*^*fl/fl;Lyz-Cre*^ (*March5* cKO) mice. All tissues except BMDMs from *March5* cKO expressed *March5* mRNA (Fig EV1B). Similarly, BMDMs derived from *March5* cKO mice did not express the MARCH5 protein (Fig EV1C).

To investigate whether MARCH5 plays an immune-modulatory function in response to bacterial infection, we challenged *March5*^*fl/fl*^ and *March5* cKO mice with a lethal dose of *Pseudomonas aeruginosa* intraperitoneally. *P. aeruginosa* is an opportunistic Gram-negative pathogen that can cause infections in humans, mainly in hospital patients, and trigger sepsis (Deng *et al*, 2016). The mice were monitored daily after *P. aeruginosa* infection, and we observed that all *March5*^*fl/fl*^ control mice died in 6 days. In contrast, 62.5% of *March5*^*fl/fl;Lyz-Cre*^ mice survived up to 10 days, showing a significant difference in survival (Fig 1A). *P. aeruginosa* infection also led to significant weight loss in mice. In just 5 days, *March5*^*fl/fl*^ mice lost 20% of their body weight and died, whereas *March5*^*fl/fl;Lyz-Cre*^ mice gained weight rapidly 4 days after *P. aeruginosa* infection (Fig 1B). Thus, it appeared that *March5* cKO mice were resistant to *P. aeruginosa* infection. Next, individual mice were sacrificed at 12 hr and 24 hr after *P. aeruginosa* injection, and the inflammatory cytokine levels in the serum, spleen homogenates and peritoneal fluid were determined using ELISA. The basal levels of inflammatory cytokines were low (< 50 pg/ml, data not shown) (Chathuranga *et al*, 2020). Upon *P. aeruginosa* infection, significant production of TNF-α and IL-6 was observed in both *March5*^*fl/fl*^ and *March5* cKO mice (Figs 1C and 1D), but the levels between these groups of mice were not different. Interestingly, however, the production of IL-1β and IL-18 after the *P. aeruginosa* challenge was different between *March5*^*fl/fl*^ and *March5* cKO mice; the *March5* cKO mice showed significantly lower levels of IL-1β (Fig 1E) and IL-18 (Fig 1F) production in three different types of samples after bacterial injection.

**Figure 1.**
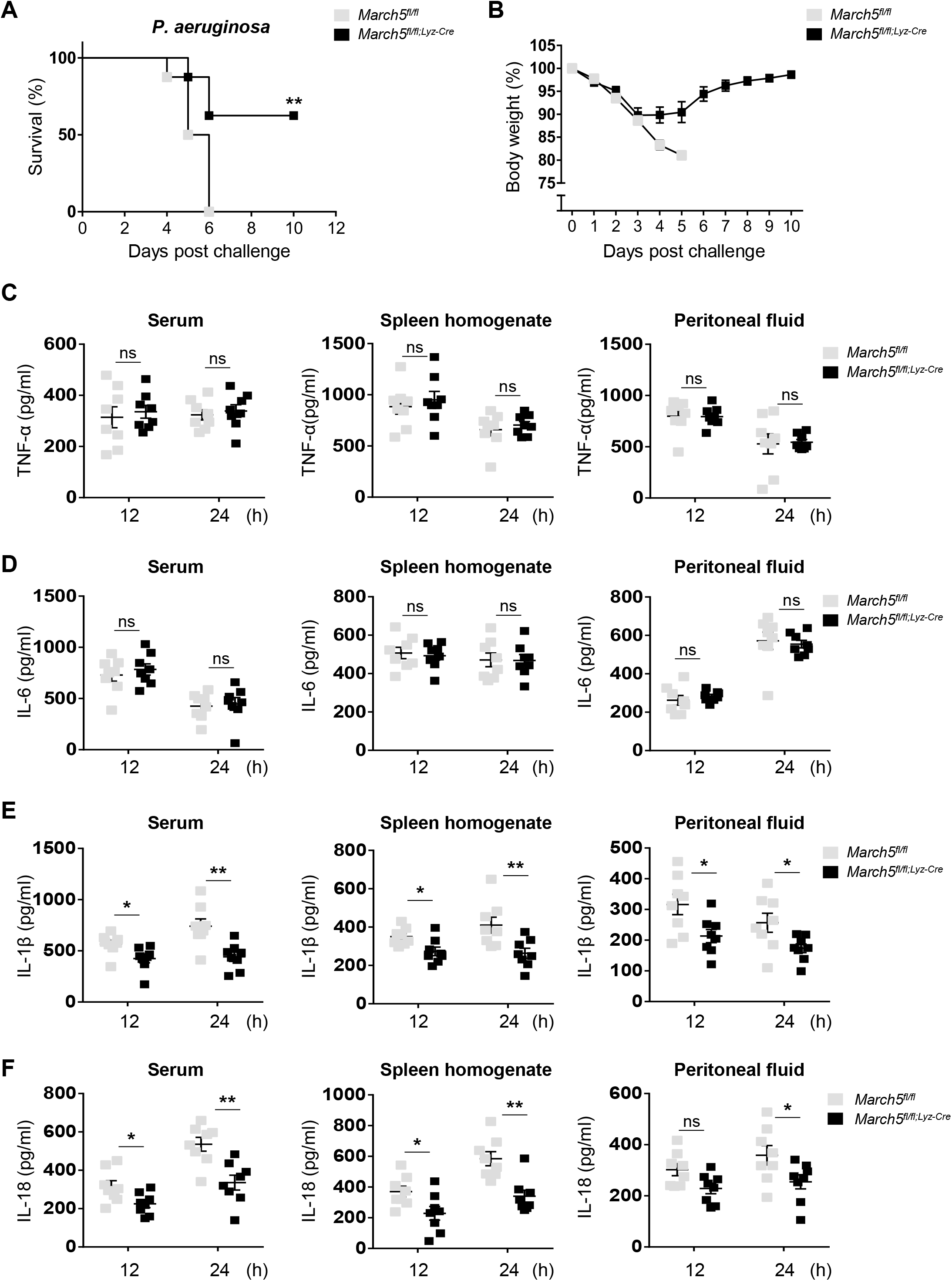
MARCH5 is required for the *in vivo* expression of IL-1β and IL-18 and lethality in response to bacterial infection. A,B *March5 cKO* mice are resistant to *P. aeruginosa* infection (A) survival rates (n = 8) and (B) changes in the body weight of *March5*^*fl/fl*^ and *March5*^*fl/fl;Lyz-Cre*^ (n = 8) mice after intraperitoneal injection with 1×10^7^ CFU *Pseudomonas aeruginosa*. C-F TNF-*α* (C), IL-6 (D), IL-1β (E) and IL-18 (F) from serum, spleen homogenate and peritoneal fluid collected from mice (n = 8) that were sacrificed 12 hr and 24 hr following the bacterial infection. Each cytokine was analyzed by ELISA. Values, *P < 0.05, **P < 0.01, ***P < 0.001 (two-tailed student’s t-test or Mantel-Cox test). Data were expressed as the mean ± SEM. See also Fig EV2.

To verify the role of MARCH5 in inflammatory response in mice, an endotoxemia-induced septic shock model was used by intraperitoneally injecting lipopolysaccharide (LPS) to mice (Raduolovic *et al*, 2018). When the mice were monitored after LPS injection (28 mg/kg body weight), 80 percent of *March5*^*fl/fl*^ control mice died 6 days after LPS injection whereas 70% of *March5* cKO mice survived up to 6 days (Fig EV2A). LPS injection caused a significant weight loss in both *March5*^*fl/fl*^ and *March5* cKO mice, showing 20% reduction in their body weight within 3 days. However, *March5* cKO mice tended to regain their body weight earlier than *March5*^*fl/fl*^ mice (Fig EV2B). Inconsistent with the observation in *P. aeruginosa* infection (Figs 1C and 1D), the ability to produce TNF-α and IL-6 in *March5*^*fl/fl*^ and *March5* cKO mice were not different after LPS injection (Figs EV2C and EV2D). However, levels of IL-1β and IL-18 after LPS challenge was different; the *March5* cKO mice showed significantly lower levels of IL-1β (Fig EV2E) and IL-18 (Fig EV2F) production. The data showed that *March5* cKO mice exhibited a significantly attenuated inflammatory response to septic shock. The data suggested that MARCH5 has a substantial impact on inflammasome activation responsible for the proteolytic processing and secretion of IL-1β and IL-18, but not the TLR4 signaling per se.

### MARCH5 potentiates NLRP3-mediated antimicrobial immunity

A diverse array of stimuli derived from microbial products and damaged host cells can induce inflammasome activation. Several cytoplasmic pattern recognition receptors recognize different types of DAMPs and PAMPs (Zheng *et al*, 2020). To identify which receptor is involved in the regulation of the inflammasome by the MARCH5 protein, BMDMs obtained from *March5*^*fl/fl*^ and *March5* cKO mice were activated with several different stimulators. To activate the NLRP3, NLRC4 and AIM2 inflammasomes in macrophages, we treated macrophages with LPS plus ATP or nigericin, flagellin (FLA-ST) and poly(dA:dT), respectively, and then measured IL-1β using ELISA and cell death by a lactic acid dehydrogenase (LDH) release assay. Cytosolic poly(I:C) has also been used to activate the NLRP3 inflammasome independently of TLR signaling. We found that treatment with LPS (200 ng/ml) alone did not change IL-1β secretion or cell death in BMDMs derived from *March5*^*fl/fl*^ mice, whereas LPS plus inflammasome activators elicited robust inflammasome activation in these cells (Figs 2A and 2B). Notably, we found that IL-1β secretion in BMDMs derived from *March5* cKO mice was significantly diminished in the context of NLRP3-mediated inflammasome signaling activation (LPS plus ATP or LPS plus Nigericin), but not NLRC4 and AIM2 inflammasome signaling activation (Fig 2A). Similarly, LDH release in BMDMs derived from *March5* cKO mice was significantly attenuated after stimulation with LPS plus ATP or LPS plus nigericin (Fig 2B), suggesting that NLRP3 inflammasome activation is specifically deficient in macrophages from *March5* cKO mice. To verify the regulatory role of MARCH5 in NLRP3 inflammasome activation, we infected BMDMs with three different bacteria (*Citrobacter rodentium, Salmonella typhimurium*, and *Pseudomonas aeruginosa*) known to stimulate the NLRP3 inflammasome (Deng *et al*., 2016; Humphries *et al*, 2018; Pu *et al*, 2017). Infection with these bacteria induced solid inflammatory responses and a substantial increase in the inflammatory cytokine release of TNF-α, IL-6, IL-1β and IL-18 in macrophages derived from *March5*^*fl/fl*^ mice (Figs 2C-E). Consistent with the finding in Fig 1, both TNF-α and IL-6 levels were comparable between *March5*^*fl/fl*^ and *March5* cKO mice (Figs 2C and 2D), whereas the production of IL-1β, IL-18 as well as caspase-1 was significantly diminished in BMDMs from *March5* cKO mice (Figs 2E and 2F, Figs EV3A-C). The inability to activate NLRP3 inflammasome by *March5* cKO mice was further confirmed by western blotting. Upon bacterial infection, *March5*^*fl/fl*^ BMDM promoted pro-caspase-1 cleavage and pro-IL-1β cleavage, whereas *March5* cKO BMDM did not (Fig EV3D). These results collectively indicated that the MARCH5 protein specifically regulates NLRP3-mediated antimicrobial immunity.

**Figure 2.**
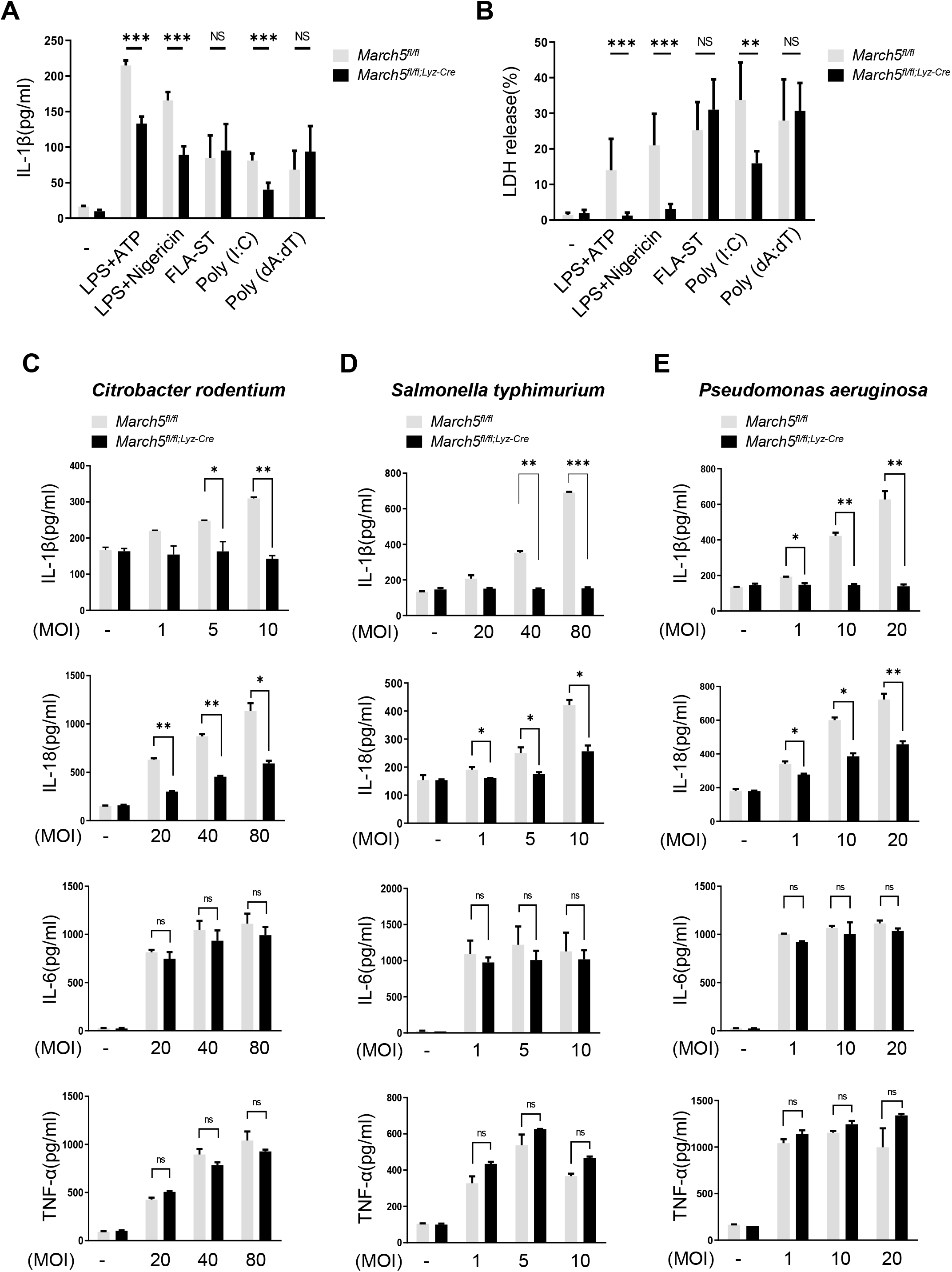
MARCH5 potentiates NLRP3-mediated antimicrobial immunity. A,B (A) IL-1β secretion and (B) LDH release were measured in the supernatants of *March5*^*fl/fl*^ and *March5*^*fl/fl;Lyz-Cre*^ BMDMs subjected to the indicated stimuli. Independent experiments were repeated at least three times, showing representative data. Values are the mean ± SD. *p < 0.1, **p < 0.01, ***p < 0.001 (two-tailed student’s t-test) C-E MARCH5 mediates activation of the NLRP3 pathway in response to bacterial infection. *March5*^*fl/fl*^ and *March5*^*fl/fl;LyzCre*^ BMDMs were infected with (C) *Citrobacter rodentium* (1 MOI, 5 MOI and 10 MOI), (D) *Salmonella typhimurium* (20 MOI, 40 MOI and 80 MOI), and (E) *Pseudomonas aeruginosa* (1 MOI, 10 MOI and 20 MOI). Secretion of IL-1β, IL-18, IL-6 and TNF-α in BMDMs infected for 12 hr was measured by ELISA. Values, *P < 0.05, **P < 0.01, ***P < 0.001 (two-tailed student’s *t-*test). Data are expressed as the mean ± SEM. See also Fig EV3.

### MARCH5 is indispensable for NLRP3 inflammasome activation

The NLRP3 inflammasome is composed of NLRP3, ASC and procaspase-1, and inflammasome activation promotes autoproteolytic cleavage of procaspase-1, the mature form of which executes the proteolytic cleavage of pro-IL-1β and pro-IL-18 (McKee & Coll, 2020). To verify whether MARCH5 is necessary for the NLRP3 inflammasome activation, we determined the cleaved IL-1β as well as caspase-1 (p20) levels in the supernatants of *March5* cKO BMDMs and MARCH5-depleted THP-1 cells after LPS primed ATP stimulation (Fig 3A, Fig EV4A). Upon treatment with LPS, the NLRP3 and pro-IL-1β levels were elevated in the cellular lysates of BMDMs and THP-1 cells. However, ATP stimulation of LPS-primed BMDMs and THP-1 cells showed a substantial difference in the levels of cleaved IL-1β and caspase-1, which were decreased in the supernatant from cultured BMDMs of *March5* cKO mice and THP-1 cells depleted of MARCH5 by siRNA. Likewise, caspase-1 activity was significantly lower in the BMDMs of *March5* cKO mice than in those of *March5*^*fl/fl*^ mice (Fig 3B). Since caspase-1 activation was deficient in macrophages from *March5* cKO mice and MARCH5-deficient THP-1 cells, we next investigated whether inflammasome assembly was affected by the absence of MARCH5 in these cells. In a coimmunoprecipitation assay using cell extracts of BMDMs stimulated with LPS and ATP, we found that the interaction between endogenous NLRP3 and ASC was weakened in macrophages of *March5* cKO mice (Fig 3C, Fig EV4B), suggesting that MARCH5 is indeed involved in NLRP3 inflammasome assembly. ASC forms oligomers in the activated inflammasome complex. To determine whether oligomerization of ASC was affected by the presence of MARCH5, cell lysates of LPS-primed BMDMs co-stimulated with nigericin were separated into Triton X-100 soluble and insoluble fractions. Triton X-100 insoluble pellets were cross-linked with DSS and subjected to western blotting. As shown in Fig 3D, ASC dimerization and oligomerization in LPS-primed *March5*^*fl/fl*^ macrophages were markedly boosted upon nigericin treatment, whereas they remained unchanged in BMDMs of *March5* cKO (Fig 3D). Similar results were observed in MARCH5-depleted THP-1 cells (Fig EV4C). The ASC speck formation assay also supported this finding. ASC assembles into a large protein complex, or “speck”, that is considered to be an upstream readout for inflammasome activation (Hoss *et al*, 2017; Stutz *et al*, 2013). Immunofluorescence staining revealed that 10∼20% of *March5*^*fl/fl*^ macrophage cells showed ASC specks after stimulation with LPS followed by ATP or nigericin and poly(dA:dT) alone. On the other hand, ASC specks were almost abolished in LPS plus ATP/nigericin stimulated BMDMs of *March5* cKO mice (Fig 3E, Fig EV4D). The number of ASC specks remained elevated in *March5* cKO BMDMs stimulated with poly(dA:dT). These data indicate that MARCH5 specifically promotes the NLRP3 inflammasome assembly and activation.

**Figure 3.**
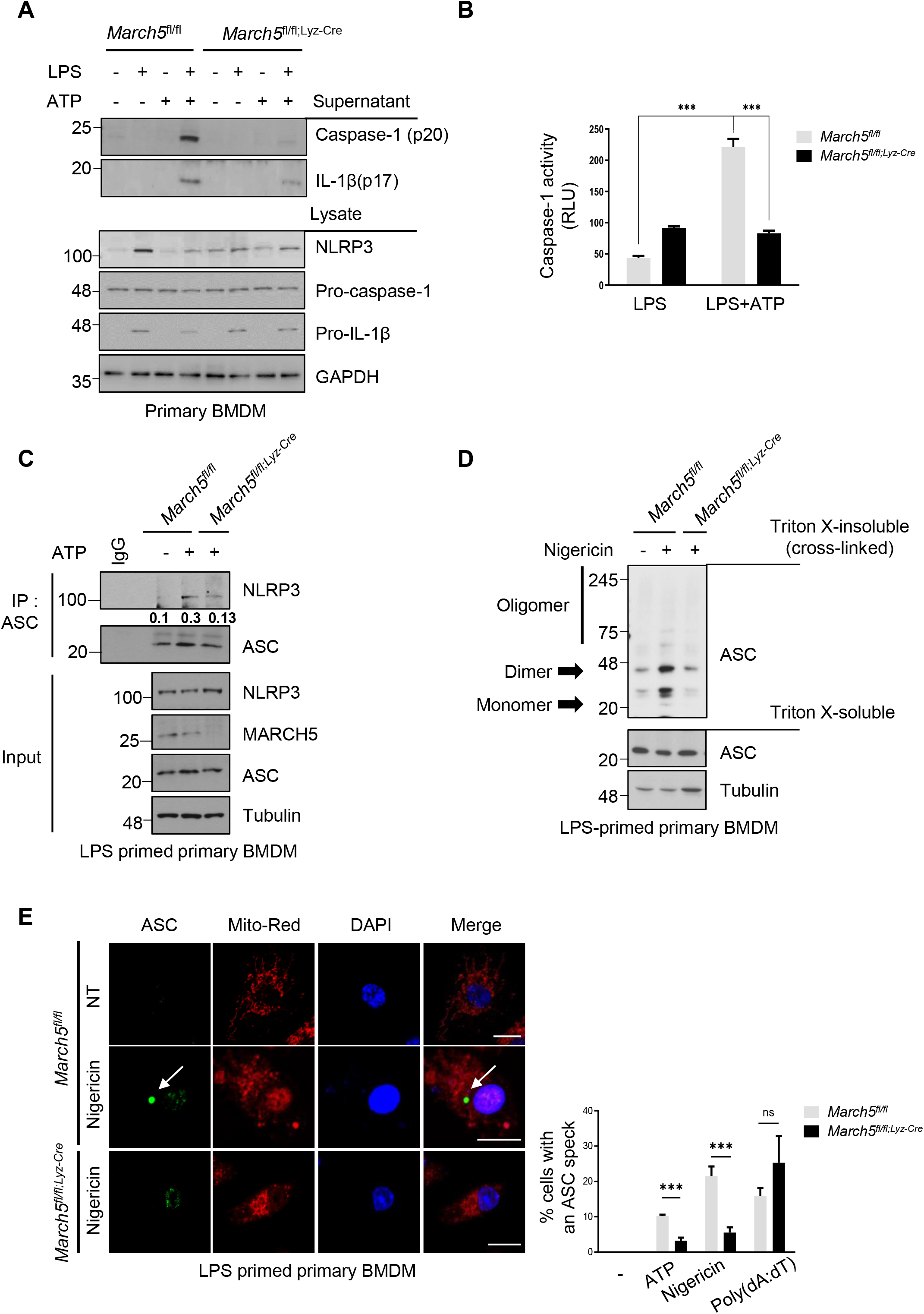
MARCH5 is essential for activating the NLRP3 inflammasome. *A March5*^*fl/fl*^ and *March5*^*fl/fl;Lyz-Cre*^ BMDMs were untreated or treated with 200 ng/ml LPS for 4 hr followed by 5 mM ATP for 45 min. The indicated proteins in the cell lysate and supernatants were immunoblotted. B The activity of caspase-1 was detected using the caspase-1 Glo assay. *March5*^*fl/fl*^ and *March5*^*fl/fl;Lyz-Cre*^ BMDMs were treated with 200 ng/ml LPS alone for 4 hr or with additional 5 mM ATP for 45 min. C,D BMDMs from *March5*^*fl/fl*^ and *March5*^*fl/fl;Lyz-cre*^ mice were primed with LPS (200 ng/ml) for 4 hr. Cells were subsequently treated with 5 mM ATP (C) or with 15µM nigericin (D). (C) Cell lysates were then immunoprecipitated with ASC antibody. Proteins were analyzed by immunoblotting with the indicated antibodies. The immunoprecipitated NLRP3 levels were quantified by using image J. (D) Triton X-100 insoluble pellets were cross-linked with 2 mM DSS and immunoblotted for ASC oligomerization. Analysis of Triton X-100 soluble and insoluble proteins was performed by immunoblotting. E *March5*^*fl/fl*^ and *March5*^*fl/fl;Lyz-cre*^ BMDMs were either transfected with 2 µg/ml poly(dA:dT) for 6 hr or treated stimulated with 200 ng/ml LPS for 4 hr along with 5 mM ATP or 15 µM nigericin for 30 min subsequently. Representative confocal images show ASC speck formation in BMDMs. The right graph shows the percentage of cells containing ASC specks. At least 100 BMDMs from each samples were analyzed. Independent experiments were repeated at least three times, showing representative data. Values are the mean ± S.D. *p < 0.1, **p < 0.01, ***p < 0.001 (two-tailed student’s t-test). Bars, 10 µm. White arrows indicate ASC speck. See also Fig EV4.

### MARCH5 interacts with the NACHT domain of NLRP3

To address the underlying mechanisms by which MARCH5 activates the NLRP3 inflammasome, we examined whether MARCH5 binds any NLRP3 inflammasome components. We transfected HEK293T cells with each Flag-tagged NLRP3 inflammasome components; Flag-NLRP3, ASC, or pro-caspase-1 along with Myc-MARCH5 and performed coimmunoprecipitation experiments with an anti-Myc antibody. Western blotting revealed that MARCH5 strongly interacted with NLRP3, coprecipitating essentially equal amounts of each inflammasome component (Fig 4A). A portion of ASC was also immunoprecipitated with MARCH5, but no pro-caspase-1 was found in this immunoprecipitant. Because overexpressed MARCH5 and NLRP3 strongly interacts in HEK293T cells, we next examined whether these interactions occurred in THP-1 cells after stimulation. In LPS-primed THP-1 cells, NLRP3 specifically interacted with endogenous MARCH5 after ATP stimulation (Fig 4B). These findings were also examined using a semi-*in vitro* immunoprecipitation assay. The LPS-primed BMDMs derived from *March5* cKO mice were co-stimulated with ATP, and cell lysates from each time point were incubated with GST-MARCH5 for the indicated durations. In the pull-down assay, MARCH5 did not bind to LPS-stimulated NLRP3 without ATP stimulation; rather, it firmly bound NLRP3 at 20 and 30 min after ATP stimulation (Fig 4C). These data suggested that MARCH5 actively interacts with NLRP3, probably in the context of the NLRP3 inflammasome. Next, we examined which domains of NLRP3 are responsible for the interaction with MARCH5. A series of deletion mutants of Myc-NLRP3 were transfected into HEK293T MARCH5 KO cells along with SFB (S-tag, Flag, and a streptavidin-binding tag)-MARCH5, and cell lysates were coimmunoprecipitated with streptavidin beads, followed by western blotting. We found that MARCH5 binding was maintained by NLRP3 mutants lacking the PYD, linker, or LRR domain, but was lost in the mutant lacking the NACHT domain (Fig 4D). Taken together, these data indicate that MARCH5 interacts with the NACHT domain of NLRP3.

**Figure 4.**
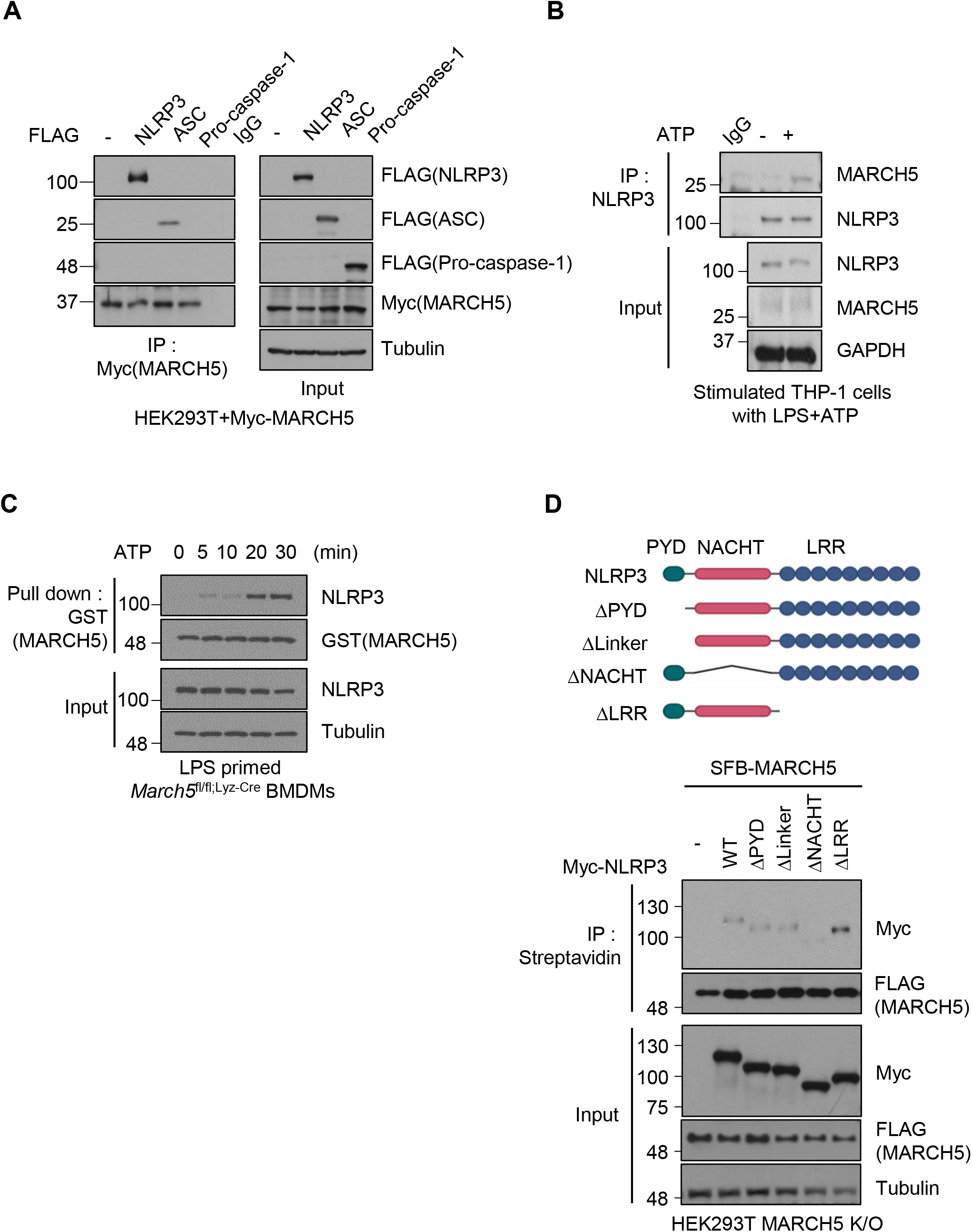
MARCH5 interacts with the NACHT domain of NLRP3. A HEK293T cells were cotransfected with Myc-MARCH5 and FLAG-tagged NLRP3, ASC, or pro-caspase-1. Cell lysates were immunoprecipitated with anti-FLAG-M2 beads and detected with the indicated antibodies. B THP-1 cells primed with 200 ng/ml LPS for 4 hr were subsequently treated with or without 5 mM ATP for 30 min. Cell lysates were immunoprecipitated with IgG or anti-NLRP3 antibody, and the indicated proteins were detected by immunoblotting. *C March5*^*fl/fl*^ BMDMs were treated with 200 ng/ml LPS for 4 hr followed by 5 mM ATP for different durations. Cell lysates mixed with GST-MARCH5 protein were pulled down with GST-tagged beads, and the indicated proteins were detected by immunoblotting. D Schematic representation of NLRP3 wild-type and truncated mutants (Upper). HEK293T cells were cotransfected with NLRP3 WT or NLRP3 mutants and SFB-MARCH5 WT. Treatment with 200 ng/ml LPS and 15 µM nigericin. Cell lysates were immunoprecipitated with streptavidin beads and immunoblotted with the indicated antibodies (Lower).

### MARCH5 promotes NLRP3 activation via K27-linked polyubiquitination of the Lys324 and Lys430 residues of NLRP3

MARCH5 maintains cellular and mitochondrial homeostasis by linking ubiquitin to target proteins (Park *et al*, 2014). To determine whether MARCH5 ubiquitinates NLRP3, we transfected FLAG-tagged NLRP3 and HA-tagged ubiquitin construct into MARCH5 WT and MARCH5 KO HEK293T cells. Immunoprecipitation with an anti-FLAG antibody revealed that the ubiquitination of NLRP3 was reduced in MARCH5 KO cells (Fig 5A). We previously showed that MARCH5 often promotes degradation of its target proteins through Lys (K) 48-linked ubiquitination (Park *et al*., 2020; Park *et al*., 2010). Thus, we determined whether MARCH5 alters the protein level of NLRP3 in the NLRP3 inflammasome complex. BMDMs obtained from *March5*^*fl/fl*^ and *March5* cKO mice were stimulated with LPS plus ATP and the protein levels of the NLRP3 inflammasome component were compared (Fig EV5A). We found that NLRP3 level was not elevated in macrophages of *March5* cKO mice, suggesting that MARCH5-dependent ubiquitination on NLRP3 does not promote the proteasome-dependent degradation pathway. Next, we carried out a ubiquitin assay after overexpression of HA-ub-K48R to exclude K48-linked polyubiquitination in MARCH5 KO HEK293T cells. We utilized the MARCH5 K40/54R(MARCH5 2KR) and MARCH5 H43W mutant constructs in this assay (Karbowski *et al*, 2007; Kim *et al*, 2016). In MARCH5 2KR, two Lys residues in MARCH5 were switched to Arg to prevent autoubiquitination and degradation of MARCH5 (Kim *et al*., 2016). The MARCH5 H43W mutant lacks the catalytic activity of MARCH5 and was used as a negative control (Karbowski *et al*., 2007). In this experiment, we found that MARCH5 2KR but not MARCH5 H43W enhanced NLRP3 polyubiquitination independent of K48-linked ubiquitin (Fig S5B). The kinetics of MARCH5-mediated NLRP3 ubiquitination during inflammasome activation was determined in cells transfected with HA-ub-K48R. NLRP3 ubiquitination by MARCH5 in LPS-primed MARCH5 KO HEK293T cells could be observed at 30 min after nigericin stimulation and became stronger at 60 min (Fig 5B).

**Figure 5.**
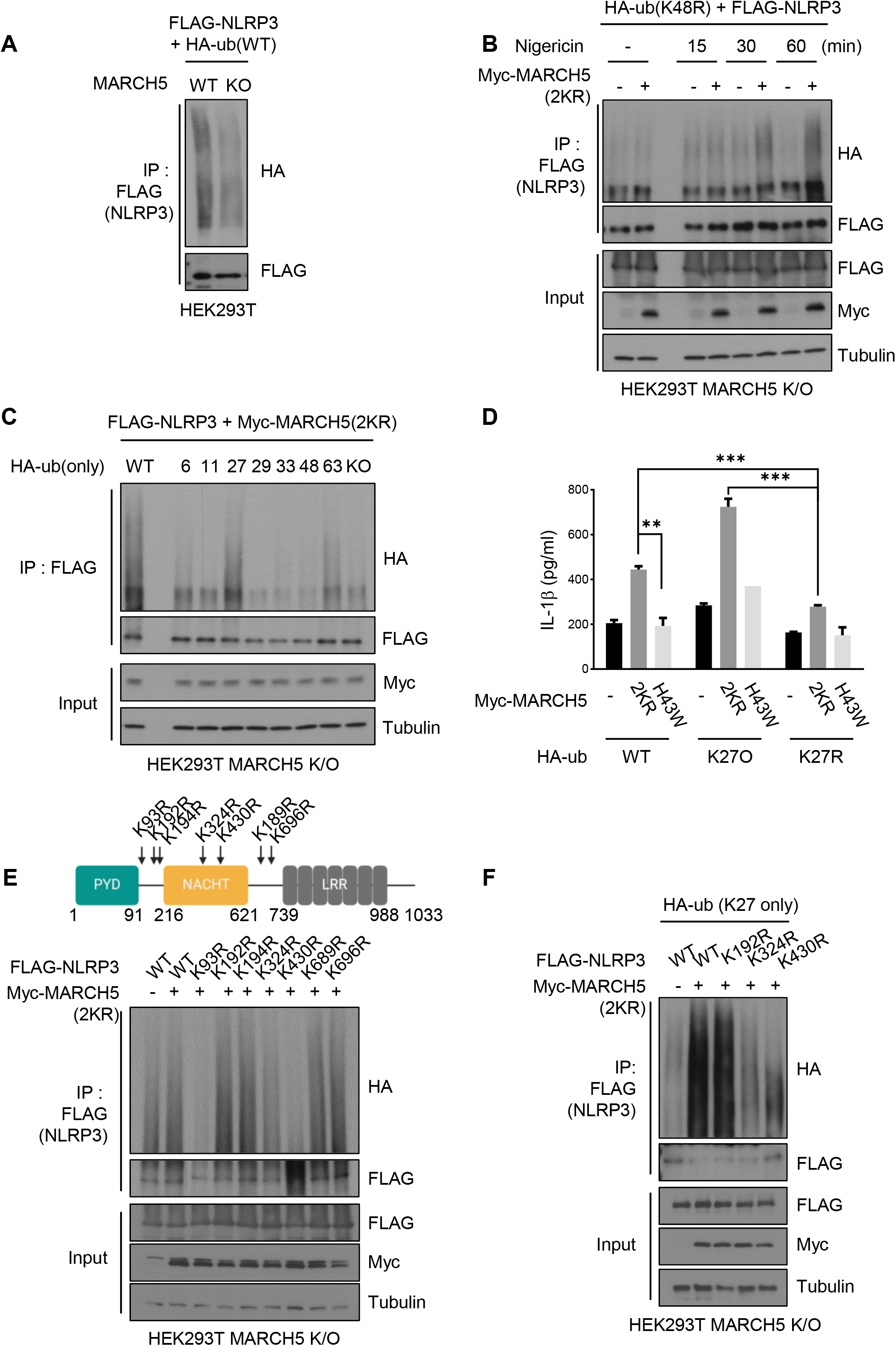
MARCH5 ubiquitinates NLRP3 on K324 and K430 via K27-linked polyubiquitination. A MARCH5 WT and KO HEK293T cells were transfected with FLAG-NLRP3 and HA-ub. After the cells were stimulated with LPS (200 ng/ml for 4 hr) and nigericin (15 µM for 60 min), the cell lysates were immunoprecipitated with FLAG-M2 beads. Ubiquitinated NLRP3 was detected by anti-HA antibody. B FLAG-NLRP3 and HA-ub K48R mutant plasmids were transfected into MARCH5 KO HEK293T cells with or without Myc-MARCH5 2KR. Then, the cells were stimulated with 200 ng/ml LPS followed by 15 µM nigericin for the indicated durations. The cell lysates were immunoprecipitated with FLAG-M2 beads and immunoblotted by using indicated antibodies. C MARCH5 KO HEK293T cells were transfected with FLAG-NLRP3, Myc-MARCH5 2KR, and each HA-ubiquitin, shown by the number of the remaining single Lys residue with the other Lys changed to Arg. For stimulation, the cells were treated with 200 ng/ml LPS for 4 hr, followed by 15 µM nigericin for 30 min. Cell lysates were immunoprecipitated with FLAG-M2 beads and analyzed by immunoblotting with the indicated antibodies. D The NLRP3 inflammasome was reconstituted in HEK293T MARCH5 KO cells expressing ASC, pro-caspase-1, and IL-1β with HA-ub WT, K27 only (K27O) mutant, or K27R mutant. Additionally, cells were cotransfected with or without Myc-MARCH5 2KR or the Myc-MARCH5 H43W mutant. Following stimulation with LPS for 4 hr and nigericin for 30 min, IL-1β secretion was quantitated by ELISA. E Schematic representation of NLRP3 Lys mutants (Upper). HEK293T MARCH5 KO cells were cotransfected with FLAG-NLRP3 WT or indicated Lys point mutants, HA-ubiquitin, and Myc-MARCH5 (2KR). Cells were stimulated with 200 ng/ml LPS for 4 hr followed by 15 µM nigericin treatment for 30 min. The cell lysates were immunoprecipitated with FLAG-M2 beads. NLRP3 ubiquitination was assessed via western blotting using anti-HA. And each protein was detected by indicated antibodies (Lower). F FLAG-NLRP3 WT or indicated Lys point mutants, HA-ub K27 only mutant and Myc-MARCH5 2KR were transfected into MARCH5 KO HEK293T cells. Cell lysates were immunoprecipitated with FLAG-M2 beads. Ubiquitinated NLRP3 was detected by immunoblotting using an HA antibody. Representative data are shown from independent experiments that were repeated at least three times. See also Fig EV5.

Next, we determined which domain of NLRP3 was ubiquitinated by MARCH5 by cotransfecting each truncated NLRP3 construct with HA-ub-K48R and MARCH5 in MARCH5 KO HEK293T cells. The data showed that the ubiquitination level of NLRP3 ΔNACHT remained unchanged upon overexpression of MARCH5. In contrast, the polyubiquitination levels of WT NLRP3, NLRP3 ΔLinker and other truncation mutants were increased by MARCH5 (Fig EV5C). Thus, we concluded that the NACHT domain of NLRP3 is the major ubiquitination target domain of MARCH5.

Next, we addressed which ubiquitination chain is mainly attached to NLRP3 among seven lysine residues (K6, K11, K27, K29, K33, K48, and K63) on ubiquitin. To do this, we utilized different ubiquitin constructs that contained only one intact Lys residue, and the other six Lys residues were mutated to Arg. We cotransfected each ubiquitin mutant with Myc-MARCH5 and FLAG-NLRP3, and immunoprecipitation with an anti-FLAG antibody showed that K27-only ubiquitin (K27O ub) was the most effective NLRP3 ubiquitination (Fig 5C). We then examined whether K27-linked polyubiquitination occurs on the NACHT domain of NLRP3 by transfecting NLRP3 truncation mutants with K27O ub. We also found that K27-linked polyubiquitination of NLRP3 ΔNACHT did not occur under the condition of MARCH5 overexpression (Fig EV5D). Thus, we concluded that MARCH5 promotes NLRP3 activation via K27-linked polyubiquitination. To investigate the physiological importance of the K27-linked polyubiquitination of NLRP3 in inflammasome activation, we compared the secretion of IL-1β in the NLRP3 inflammasome reconstitution system after cotransfection with K27O ub or K27R ub. Indeed, a significant increase in IL-1β secretion was found in cells transfected with K27O ub and MARCH5, whereas this increase was not observed in cells transfected with K27R ub or MARCH5 H43W, although the expression of the transfected proteins was comparable in this system (Figs 5D and EV5E). Taken together, these results indicate that MARCH5 activates the NLRP3 inflammasome via K27-linked polyubiquitination of the NACHT domain of NLRP3.

We next addressed which Lys residues of NLRP3 are ubiquitinated by searching for putative ubiquitination sites on NLRP3 using the UbPred program. This analysis predicted seven ubiquitination sites on Lys (K9, K192, K194, K324, K430, K689, K696) of NLRP3. Based on this analysis, we generated 7 Lys point mutants in which each Lys was replaced with Arg (Fig 5E, upper). We cotransfected each FLAG-tagged NLRP3 Lys mutant with the MARCH5 2KR and HA-ub constructs in MARCH5 KO HEK293T cells, and after cotreatment with LPS plus nigericin, polyubiquitination was examined in the immunoprecipitant obtained using anti-FLAG beads. We found that polyubiquitination of NLRP3 was significantly reduced in the cells transfected with K93R, K324R and K430R mutants, whereas other Lys mutants of K192R, K194R, K689R, and K696R of NLRP3 showed polyubiquitination levels equivalent to those of the WT (Fig 5E, lower). Since MARCH5 targeted the NLRP3 NACHT domain, we focused on the two Lys sites that were in the NACHT domain, K324 and K430. To further confirm the lysine sites on NLRP3 targeted by MARCH5 for K27-linked polyubiquitination, we carried out a ubiquitination assay after transfection with K27O ub, where the K192R mutant of NLRP3 was used as a positive control. Consistent with Fig 5E, K27-linked polyubiquitination of NLRP3 was reduced in the K324 and K430 mutants of NLRP3 (Fig 5F). Our data suggested that K27-linked polyubiquitination by MARCH5 occurred at the K324 and K430 sites on NLRP3.

### MARCH5-dependent NLRP3 ubiquitination allows NEK7 binding to NLRP3

To address the functional role of NLRP3 ubiquitination by MARCH5, we examined NLRP3 oligomerization through fluorescence microscopy. Unlike the ASC specks, the NLRP3 puncta were barely visible, probably due to prompt inflammasome assembly with ASC and pro-caspase-1. We utilized immortalized BMDM (iBMDM) cells expressing GFP-tagged NLRP3 and knocked down ASC by siRNA. Upon treatment with LPS plus nigericin, oligomerized NLRP3 puncta were found in ∼ 20% of cells, but these puncta were significantly diminished in *March5*-depleted iBMDMs (Figs 6A and EV6A). In SDD–AGE (semi-denaturing detergent agarose gel electrophoresis), the high-molecular weight oligomers of NLRP3 was diminished in *March5* cKO BMDMs stimulated with LPS and nigericin or ATP at two different time points (Fig 6B), suggesting that MARCH5-mediated ubiquitination facilitates NLRP3 self-oligomerization. MAVS recruits NLRP3 to the mitochondria for inflammasome activation (Park *et al*, 2013; Subramanian *et al*., 2013). It also shows that NEK7 directly binds NLRP3, and these interactions are essential for NLRP3 oligomerization (He *et al*, 2016; Sharif *et al*., 2019). As shown in Fig EV6B, coimmunoprecipitation with anti-NLRP3 antibody revealed that NLRP3 interacts with MAVS in BMDMs treated with LPS plus nigericin, and the absence of MARCH5 did not disrupt these interactions. Consistently, ubiquitination-defective NLRP3 K324R and K430R mutants remained bound to MAVS (Fig EV6C). On the contrary, depletion of MAVS by siRNA disrupted the interaction of MARCH5 with NLRP3, which furthermore disrupted the interaction of ASC with NLRP3 (Fig EV6D). Thus, the data suggest that MAVS mediates the mitochondrial recruitment of NLRP3 that is subsequently ubiquitinated by MARCH5, which is essential for the inflammasome assembly. Next, we addressed whether MARCH5 affects NEK7 binding to NLRP3. We knocked down MARCH5 (MARCH5 KD) in THP-1 cells using siRNA, followed by stimulation with LPS and ATP. Coimmunoprecipitation with an anti-NLRP3 antibody revealed that the interaction between NLRP3 and NEK7 was impaired in these cells. Accordingly, the association of NLRP3 with ASC was also diminished (Fig 6C). To examine the multimeric complex formation of NLRP3 oligomers with NEK7, stimulated iBMDM cell lysates were separated on a blue native gel, followed by SDS−PAGE in a second dimension. As expected, NLRP3 and NEK7 were found in a high molecular weight region (Fig 6D, yellow circle). In contrast, these signals were not present in lysates from MARCH5 KD iBMDM cells, suggesting that MARCH5 is indispensable for forming NEK7-NLRP3 multimeric oligomers. Indeed, wild-type NLRP3 bound endogenous NEK7 in HEK293T cells, whereas the binding of NLRP3-K324R and NLRP3-K430R mutants to NEK7 was strongly reduced (Fig 6E). In consistent with previous reports, the NLRP3-S198A mutant, which is defective in JNK1-dependent phosphorylation (Song *et al*, 2017), also barely binds NEK7. In addition, the ability to form multimeric oligomers of NEK7-NLRP3 was impaired in cells with NLRP3-K324R and NLRP3-K430R mutants, as shown in a two-dimensional polyacrylamide gel electrophoresis (2D-PAGE) (Fig 6F). Finally, the colocalization of NEK7 with NLRP3 puncta was significantly reduced, as shown by immunofluorescence staining (Fig 6G). Together, these data indicate that MARCH5-dependent NLRP3 ubiquitination is necessary for the binding of NEK7 and the formation of NEK7-NLRP3 oligomers.

**Figure 6.**
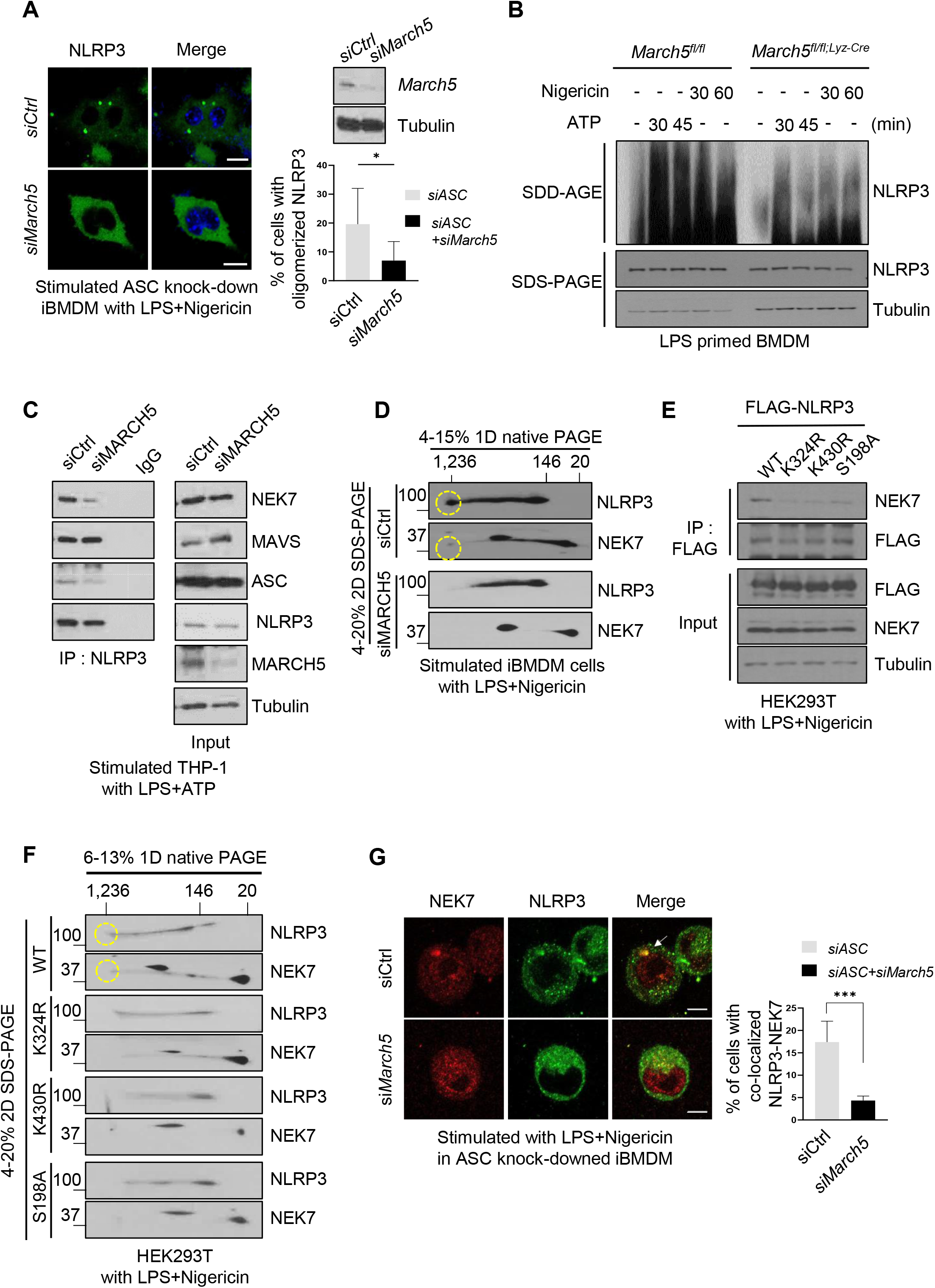
MARCH5-dependent NLRP3 ubiquitination allows the binding of NEK7 to NLRP3. A Representative confocal images of oligomerized NLRP3. ASC knockdowned immortalized BMDMs expressing GFP-tagged NLRP3 WT were transfected using indicated siRNAs, and treated with 200 ng/ml LPS for 4 hr followed by 15 µM nigericin for 30 min. White arrows indicate oligomerized NLRP3. The percentage of cells with oligomerized NLRP3 was quantified. At least 100 cells were analyzed. Bar, 10 µm. Values are the mean ± SD. * p < 0.1, ** p < 0.01, *** p < 0.001 (two-tailed student’s t-test). *B March5*^*fl/fl*^ and *March5*^*fl/fl*;Lyz-Cre^ BMDMs were stimulated for 4 hr with LPS followed by 5 mM ATP or 15 µM nigericin for the indicated durations. SDD–AGE and SDS−PAGE were performed to separate cell lysates, followed by immunoblotting using indicated antibodies. C THP-1 cells transfected with siControl or siMARCH5 were stimulated with 200 ng/ml LPS for 4 hr followed by 5 mM ATP for 30 min. Cell lysates were subjected to immunoprecipitation by using NLRP3 antibody. Western blotting was performed using the indicated antibodies. D Immortalized BMDMs were transfected with siControl or siMARCH5 and stimulated with LPS and nigericin. Blue native PAGE was used first to separate the cell lysates, and then SDS−PAGE was used to separate them further. Western blotting was performed using the indicated antibodies. E HEK293T cells were transfected with FLAG-tagged NLRP3 WT or indicated mutants and stimulated with LPS and nigericin. Cell lysates were immunoprecipitated with FLAG-M2 beads and detected with the indicated antibodies. F HEK293T cells were transfected with FLAG-tagged NLRP3 WT or indicated mutants. Cells were stimulated with LPS and nigericin. Cell lysates were separated by blue-native PAGE (first dimension) and SDS−PAGE (second dimension). Representative data from independent experiments that were repeated at least three times are shown. G Representative confocal image showing NLRP3-NEK7 colocalization. siControl-and si*March5-* transfected ASC knockdowned iBMDMs expressing GFP-tagged NLRP3 WT cells were stimulated with 200 ng/ml LPS for 4 hr followed by 15 µM nigericin for 30 min. The percentage of cells with NLRP3-NEK7 colocalization was quantified. At least 100 cells were analyzed. Values are shown as the mean±SD. *p < 0.1, **p < 0.01, ***p < 0.001 (two-tailed student’s t-test). See also Fig EV6.

### MARCH5-mediated ubiquitination of NLRP3 is an essential step for inflammasome activation

Next, we examined the physiological relevance of these two Lys residues of NLRP3 by assessing their roles in NLRP3 inflammasome activation. We transfected ASC with NLRP3 WT or NLRP3 Lys mutant in HEK293T cells and treated them with nigericin. As shown in Fig 7A, approximately 30% of the cells expressing WT NLRP3 formed ASC specks, while cells expressing NLRP3-K324R and NLRP3-K430R showed less ASC speck. Consistent with this result, we confirmed ASC oligomerization by SDD-AGE. We overexpressed the NLRP3-K324R and NLRP3-K430R constructs with ASC in HEK293T cells. After treatment with LPS plus nigericin, cell lysates were treated with DSS and divided into a Triton X-100 soluble fraction and Triton X-100 insoluble pellet. Immunoblotting showed that NLRP3-WT and NLRP3-K93R were able to form ASC oligomers (Fig 7B). In contrast, NLRP3-K324R and NLRP3-K430R failed to stimulate ASC oligomerization, similar to NLRP3-S198A. In the reconstituted NLRP3 inflammasome complex, the mature form of IL-1*β* was reduced in the cells transfected with NLRP3-K324R or NLRP3-K430R, unlike those transfected with NLRP3-WT and NLRP3-K93R, which exhibited a normal level of mature IL-1β (Fig 7C). The NLRP3-S198A mutant also formed a nonfunctional NLRP3 inflammasome, as expected. Similarly, IL-1*β* secretion was substantially diminished in cells expressing NLRP3-K324R, NLRP3-K430R, or NLRP3-S198A (Fig 7D). These results suggest that MARCH5-mediated ubiquitination of the Lys324 and Lys430 residues of NLRP3 is an important event in NLRP3 inflammasome assembly and activation.

**Figure 7.**
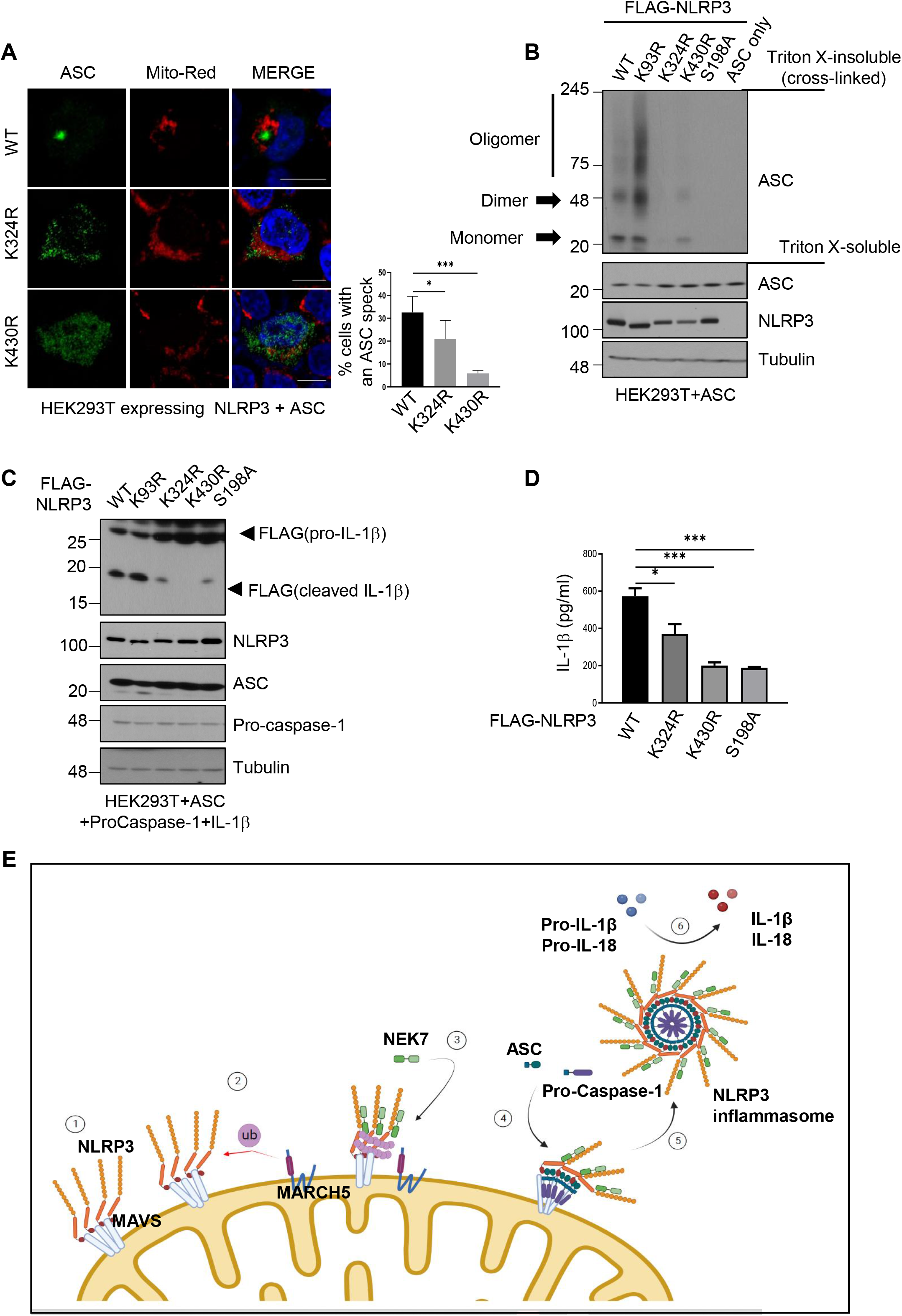
MARCH5-mediated ubiquitination of NLRP3 is essential for inflammasome activation. A HEK293T cells were transfected with NLRP3 WT or indicated Lys mutants and ASC. Cells were stimulated with 200 ng/ml LPS for 4 hr followed by 15 µM nigericin for 30 min. The percentage of cells with an ASC speck was quantified after Confocal microscopy. At least 100 cells were analyzed. Values are the mean ± SD. Values, *p < 0.1, **p < 0.01, ***p < 0.001 (two-tailed student’s t-test). B HEK293T cells were transfected with ASC and NLRP3 (WT or individual NLRP3 mutants). Cells were treated for 4 hr with 200 ng/ml LPS and 30 min with 15 µM nigericin. Harvested cell lysates were incubated with 2 mM DSS for cross-linking. Triton X-100 insoluble pellets and soluble fraction lysates were detected by immunoblotting with the indicated antibodies. C NLRP3 inflammasome-reconstituted HEK293T cells with NLRP3 WT or indicated mutants were stimulated for 4 hr with 200 ng/ml LPS and 30 min with 15 µM nigericin. Harvested cells were detected by immunoblotting with the indicated antibodies. D Culture supernatants were obtained from (C) and subjected to ELISA to quantify secreted IL-1β. Representative data are shown from independent experiments that were repeated at least three times. Values are the mean ± SD. *p < 0.1, ** p <0.01, *** p <0.001 (two-tailed student’s t-test) E Schematic diagram of NLRP3 inflammasome activation by MARCH5. Upon activation by a priming signal, NLRP3 associates with MAVS on mitochondria ①. MARCH5 recognizes and ubiquitinates two lysine sites, K324 and K430, on the NLRP3 NACHT domain via K27-linked polyubiquitination ②. NLRP3 ubiquitination by MARCH5 promotes the recruitment of NEK7 and enables self-oligomerization ③. As a result, ASC and pro-caspase-1 are recruited to oligomerized NLRP3 ④ for NLRP3 inflammasome assembly ⑤ and trigger the activation of caspase-1 and pro-cytokine cleavage ⑥.

## Discussion

NLRP3 responds to various stimuli, and aberrant NLRP3 activation underlies autoimmune and degenerative diseases such as atherosclerosis, Alzheimer’s disease, type 2 diabetes and lupus (Fusco *et al*., 2020). A multilayered regulatory mechanism ensures accurate NLRP3 inflammasome activation, which has the benefit of presenting multiple specific molecular targets for disease therapy (McKee & Coll, 2020). In the present study, we uncovered the sequence of steps of NLRP3 assembly on the mitochondrial outer membrane (Fig 7E). When the activation signal is received, primed NLRP3 moves to mitochondria and binds MAVS (①). The mitochondrial-resident E3 ligase MARCH5 delivers ubiquitin to the NACHT domain of NLRP3 through K27-linked polyubiquitination (②). This modification triggers NEK7 binding to ubiquitinated NLRP3, forming the NEK7-NLRP3 multimeric complex (③,④). Consequently, ASC and pro-caspase-1 associate with NLRP3 oligomers to complete inflammasome assembly (⑤). Finally, the pro-inflammatory cytokines IL-1β and IL-18 are cleaved by activated caspase-1 (⑥). Thus, our data show that mitochondrial-resident MARCH5 E3 ligase is an essential regulator that initiates NEK7 binding to NLRP3. It also highlights the mitochondria as an onset platform for NLRP3 immune complex formation.

NLRP3 has been observed in the cytosol as well as in membranous organelles (the ER, mitochondria, and Golgi) during inflammasome activation (Hamilton & Anand, 2019). In resting state cells, the inactive state of NLRP3 could be monomeric or oligomeric double-ring cages embedded in the organelle membrane such that its pyrin domains are hidden inside (Andreeva *et al*, 2021). NLRP3 undergoes dynamic relocation in cells for inflammasome activation (Wang *et al*, 2013), and several activation steps are necessary for inflammasome assembly and activation (Paik *et al*, 2021). MAVS mediates NLRP3 recruitment to mitochondria through direct interaction, and this step is indispensable for NLRP3 inflammasome activation (Park *et al*., 2013; Subramanian *et al*., 2013). Indeed, the absence of MAVS in stimulated THP-1 cells disturbed the interaction of NLRP3 with ASC and abolished NLRP3 binding to MARCH5 (Fig EV6D). MARCH5-dependent ubiquitination of NLRP3 did not affect the MAVS-NLRP3 interaction, but it severely damaged its interaction with NEK7 (Fig 6E). Thus, the data suggest that NLRR3 recruitment to mitochondria by MAVS precedes its interaction with MARCH5. NLRP3 phosphorylation at Ser194 is a key priming event during LPS stimulation (Song *et al*., 2017). We observed that cells expressing the NLRP3-S194A mutant also failed to form the multimeric NEK7-NLRP3 complex (Figs 6E and F). Thus, at least two preceding events may occur before MARCH5 ubiquitinates NLRP3: phosphorylation of NLRP3 by JNK1 and mitochondrial localization of NLRP3 by MAVS. In ASC knock down cells, fluorescence microscopy using GFP-NLRP3 revealed oligomerized NLRP3 puncta upon stimulation with LPS plus nigericin (Figs 6A and EV6A). ASC specks are limited to one per cell near the MTOC, but more NLRP3 puncta per cell can form in ASC KD cells.

Interestingly, microtubule affinity regulating kinase 4 (MARK4) interacts with NLRP3 and facilitates NLRP3 localization to the mitochondria. MARK4 promotes the relocation of NLRP3 to the MTOC in a microtubule-dependent manner (Li *et al*, 2017). Thus, our data suggest that mitochondria serve as an onset platform for NLRP3-NEK7 complex formation and that subsequent association with ASC and pro-caspase-1 occurs near the MTOC. Recently, it has been reported that NLRP3 recruited to Golgi complex did not require NEK7 for its activation (Schmacke *et al*, 2022). Thus, there are, at least, two main pathways for NLRP3 activation in cells; NEK7-dependent and -independent NLRP3 activation in mitochondria and Golgi apparatus, respectively.

NLRP3 forms an autoinhibitory conformation in a resting state and undergoes a conformational change upon inflammasome stimulation. Self-oligomeric assembly of NLRP3 requires its NACHT domain, which directly interacts with the neighboring NLRP3. The NACHT domain comprises 4 subdomains, NBD, HD1, WHD (winged helix domain), and HD2, and a drastic conformational rearrangement among these subdomains occurs, positioning the NLRP3 NBD-HD1 module to a direct interacting interface between neighboring NLRP3 molecules (Sharif *et al*., 2019). Notably, NEK7 is a crucial upstream mediator of NLRP3 oligomerization. Among the NLR receptors, NEK7 specifically binds NLRP3 (He *et al*., 2016). A cryo-EM structure study revealed that NEK7 binds to the LRR domain and NACHT domain (HD2) of inactive NLRP3 (Sharif *et al*., 2019). An as-yet unidentified priming or activation step is required to induce rotational activation of the NACHT domain. This rotational activation step is necessary for uncovering part of the NACHT surface, enabling NLRP3 oligomerization. We demonstrated that MARCH5 ubiquitinates the NACHT domain of NLRP3 and targets two lysine residues (K324 and K430) of NLRP3 in the NBD and HD1 subdomains, respectively. Multimeric NEK7-NLRP3 complex formation was abrogated in cells depleted of MARCH5 (Fig 6D) or cells expressing NLRP3 mutants that are defective for ubiquitination at these lysine residues (Fig 6F). Under these conditions, the NLRP3-NEK7 interaction was impaired (Figs 6C and E). Thus, it can be proposed that upon inflammasome stimulation, NLRP3 attaches to MAVS on the mitochondrial outer membrane, and MARCH5-mediated ubiquitination of the NLRP3 NACHT domain might trigger its rotational switch to allow NEK7 to bind NLRP3, forming a large oligomeric complex of NEK7 and NLRP3.

Taken together, our data show that MARCH5 plays a key role in regulating NLRP3 inflammasome activation. The mechanism of MARCH5-mediated NLRP3 inflammasome activation provides new insight into the NEK7-NLRP3 binding signal and NLRP3 assembly for inflammasome activation on the mitochondria. Uncontrolled NLRP3 inflammasome activation is involved in several human diseases. Although blocking the cytokines or molecules downstream of the inflammasome is a common strategy for treating several diseases, cytokine secretion and pyroptosis by inflammasome activation are both pathogenic. Therefore, targeting inflammasome assembly would be more effective therapy for these diseases. Targeting the interaction of NLRP3 and MARCH5 or regulating MARCH5 activity may represent new therapeutic strategies for NLRP3-associated inflammatory disease.

## Materials and Methods

### Mice

*March5*^*tm1a*^ mice on a C57BL/6 background harboring LoxP sites flanking *March5* exon3 were purchased from the European Mouse Mutant Archive. We cross-bred Lysozyme M-Cre mice to generate *March5* cKO mice (*March5*^*tm1c*^). All mice were maintained in a specific pathogen-free animal facility. The genotypes of *March5* animals were determined by PCR using a specific primer: forward 5’-GTGACACTACTTTTGATGTGAAG-3’, reverse 5’-ATGCTACAGCTCATGTGTAAG-3’. For the mice experiment, 12∼20 weeks old male mice were used. Chungnam National University’s Institutional Animal Use and Care Committee approved all animal experiments (CNU-00777, 202109A-CNU-168). They were performed in biosafety level BSL-2 laboratory facilities with the Guide for the Care and Use of Laboratory Animals (published by the US National Institutes of Health).

### LPS septic shock model

LPS (28 mg/kg body weight) was given to mice aged 6-8 weeks via intraperitoneal injection. Serum, spleen and peritoneal lavage were obtained after 12hr and 24 hr, and cytokine levels were measured using ELISA. For survival analysis, mice were administered LPS (28 mg/kg body weight) intraperitoneally and monitored for 8 days.

### *P. aeruginosa* infection

Six to eight weeks old *March5*^*fl/fl*^ and *March5* cKO mice were inoculated intraperitoneally with *P. aeruginosa* (ATCC, BAA-1744) (100 µl; 1 × 10^7^ CFU) suspended in sterile endotoxin-free PBS. The animals were sacrificed at 12 hr and 24 hr post-inoculation, and blood, spleen, peritoneal lavage were collected. Coagulated blood was centrifuged at 16,000 × g for 15 min, and the supernatants were collected as serum. Spleen homogenate, serum and peritoneal fluid samples were assayed for TNF-*α*, IL-6, IL-1β and IL-18. For survival analysis, mice were administered with *P. aeruginosa* (100 µl; 1 × 10^7^ CFU) intraperitoneally and monitored for 10 days.

### Cell Culture

THP-1 cells were cultured in RPMI1640 with 10% heat-inactivated FBS and 1% antibiotic-antimycotic. For differentiation, THP-1 cells were treated with 100 nM Phorbol-12-myristate-13-acetate (PMA) for 72 hr. Bone marrow-derived macrophages (BMDMs) were isolated from femurs and tibias of 12∼20 weeks-old mice. After collection of bone marrow, red blood cells were removed by ACK lysis buffer (GIBCO BRL), and cells were cultured in RPMI1640 supplemented with 10% heat-inactivated FBS and 1% antibiotic-antimycotic containing 20% L929 cell-conditioned medium. HEK293T (ATCC, ACS-4500) cells were cultured in DMEM supplemented with 10% heat-inactivated FBS and 1% antibiotic-antimycotic. Immortalized BMDMs (iBMDMs) were kindly provided by Dr. Je-Wook Yu and cultured in DMEM with 10% heat-inactivated FBS and 1% antibiotic-antimycotic with 2% L929 cell-conditioned medium. All cells were cultured at 37°C in 5% CO_2_ incubator.

### Western blotting

Cells were lysed in SDS sample buffer and boiled for 10 min. The cell lysate was separated by 4-20% gradient SDS-PAGE and transferred to the nitrocellulose membrane. Immunoblots were detected by ECL system (GE Healthcare) and analyzed by using the following antibodies : anti-NLRP3 (1:10,000, AG-20B-0014, Adipogen), anti-NLRP3 (1:1,000, #15101S, Cell signalling), anti-mouse ASC (1:1,000 AG-25B-0006, Adipogen), anti-human ASC (1:1,000, sc-22514-R, Santa Cruz biotechnology), anti-ASC (1:1,000, sc-514414, Santa Cruz biotechnology), anti-human IL-1β (1:1,000, #12242, Cell Signaling Technology), anti-IL-1β (1:1,000, sc-12742, Santa Cruz biotechnology), anti-mouse caspase-1 (1:1,000, AG-20B-0042, Adipogen), anti-human caspase-1 (1:1,000, #3866, Cell Signaling Technology), anti-caspase-1 (1:1,000, sc-56036, Cell signalling technology,), anti-mouse IL-1β (1:1,000, AF-401-NA, R&D system), anti-human MAVS (1:1,000. #3993, Cell signaling technology), anti-FLAG (1:10,000, F1804, Sigma Aldrich), anti-c-Myc (9E10) (1:1,000, sc-40, Santa Cruz biotechnology) anti-Tubulin (1:10,000, sc-73232, Santa cruz technology), anti-GAPDH (1:5,000, #2188, Cell signaling technology), anti-HA (1:1,000, sc-7392, Santa cruz technology), anti-Ub (1:1,000, sc-8017, Santa cruz technology). To detect the release of caspase-1 (p20) and IL-1β (p18), a conditioned medium was collected and incubated with the same volume of methanol and 1/4 volume of chloroform. The mixture was centrifuged at 16,000 × g for 10 min at RT. The top layer (500µl) was removed and mixed with 1 ml of methanol by vortexing. Then the mixture was centrifuged at 16,000 × g for 10 min at RT. The supernatant was removed and dried at RT for 15 min. The protein pellet was dissolved in 2X SDS sample buffer and boiled at 100°C for 15 min, and subjected to SDS-PAGE.

### Reconstitution of NLRP3 inflammasome in HEK293T cells

HEK293T cells were cultured in 12-well plates and transfected by polyethyleneimine (PEI, Polysciences) with following constructs: FLAG-NLRP3 or Flag-NLRP3 K mutants (100 ng/well) or Myc-NLRP3(10 ng/well), Flag-ASC (10 ng/well), Flag-pro-caspase-1 (50 ng/well), pro-IL-1β-Flag (400 ng/well). After 24∼36 hr, cells were lysed with 1X sample buffer and sonicated. The expression of the protein was assessed by immunoblotting.

### Immunoprecipitation

Cells were lysed in lysis buffer (50 mM Tris, pH 7.8, 50 mM NaCl, 1% NP-40, 5 mM EDTA, 10% Glycerol, 1 mg/ml Aprotinin, 1 mg/ml Leupeptin, 5 mM NaF, 0.5 mM Na_3_VO_4_). Lysates were sonicated and centrifuged at 16,000 × g at 4°C for 20 min. The supernatant was incubated with either anti-NLRP3 or anti-Flag-M2 beads at 4°C overnight with agitation. Next day, samples were incubated more with protein A/G Sepharose beads (GE healthcare) for 2 hr washed with lysis buffer, and then eluted with 2X SDS sample buffer. Samples were separated by SDS-PAGE and subjected to immunoblotting.

### RNA interference

To knock down the protein expression in THP-1 and iBMDMs, cells were transfected with small interfering RNA (siRNA) oligonucleotides. Following siRNA were custom synthesized by Bioneer; Human-specific MARCH5 siRNA(5’-GGGUGGAAUUGCGUUUGUUTT-3’), mouse-specific March5 siRNA(5’-GGU UGU AGG CCA UAA AGA A-3’), mouse-specific ASC siRNA (5’-GCUCUUCAGUUUCACACCA-3’), and Human-specific MAVS siRNA was obtained from Bioneer(57506). siRNA was transfected in THP-1 cells by Lipofectamine 2000(Invitrogen).

### *In vivo* ubiquitination assay

HEK293T cells were transfected with HA-ubiquitin, FLAG, or Myc-tagged constructs by using polyethylenimine (PEI, Polysciences, 23966). After 24 hr later, cells were activated by 200 ng/ml LPS for 4 hr followed by 15 µM nigericin for 30∼60 min. Cell lysates were immunoprecipitated with Flag-M2 bead (Sigma-Aldrich, A2220) or anti-Myc antibody at 4 °C overnight with agitation. Beads were washed with lysis buffer four times. They were eluted with 2X SDS sample buffer, and subjected to western blotting.

### Inflammasome activation

Primary BMDMs or THP-1 cells were seeded in the culture dishes for overnight. Cells were primed for 4 hr with 200 ng/ml LPS (Sigma-Aldrich, L9274) and then stimulated as follows: 5 mM ATP (Sigma-Aldrich, A6419) for 30 ∼ 45 min, 15 µM nigericin (Invivogen, tlrl-nig) for 30 min ∼ 1hr respectively. Cells were stimulated with 250 ng/ml FLA-ST (Invivogen, tlrl-stfla) for 4 hr. Cells were transfected with 2 µg/ml poly(I:C) (Invivogen, tlrl-pic) or 2 µg/ml poly(dA:dT) (Invivogen, tlrl-patn-1) by using Lipofectamine 2000 for 6 hr. *Citrobacter rodentium* (ATCC, 51459), *Pseudomonas aeruginosa* (ATCC, BAA-1744), and *Salmonella typhimurium* strain (ATCC, 14028) grown in lysogeny broth in a log phase were resuspended in phosphate-buffered saline. For the bacterial infection, each bacteria titer was examined in serial dilution inoculating LB Agar plates. Primary BMDMs were seeded (1 × 10^6^ cells/ well in 12-well culture plates and allowed to rest for 12 hr. Cells were then stimulated with the indicated bacteria.

### ELISA

Cells were seeded into 12-well plates and treated with indicated activators and indicated bacteria. The supernatant was collected at the indicated time points and then quantified by using following commercially available ELISA kits; mouse IL-1β (Biolegend, 432616), Caspase-1 (Novus biologicals, NBP2-75014), mouse TNF-α (BD OptEIA, 555268), mouse IL-6 (BD OptEIA, 555240) and mouse IL-18 (MBL, 065FA).

### Cytotoxicity assay

BMDMs were seeded into 12-well plates and treated with activators. Lactate dehydrogenase (LDH) release to the culture supernatant was quantified by using the LDH cytotoxicity assay kit (Promega).

### ASC speck staining

Cells were primed with 200 ng/ml LPS for 4 hr, and subsequently stimulated with 5 mM ATP or 15 µM nigericin for 30 min. poly(dA:dT) were transfected using Lipofectamine 2000. After stimulation, cells were fixed with 4% paraformaldehyde for 30 min at 37°C and permeabilized with 0.1% Triton X-100 for 10 min. The slides were blocked with 2% BSA in PBS. Cells were stained with anti-ASC(1:200) and anti-NLRP3(1:300). To stain nuclei, cells were stained with DAPI. Cells were visualized by confocal microscopy (Zeiss LSM 710) at the Three-dimensional immune system imaging core facility of Ajou University.

### ASC oligomer cross-linking

Cells were primed with 200 ng/ml LPS for 4 hr, and subsequently stimulated with 5 mM ATP or 15 µM nigericin. Cells were lysed with Triton buffer (50 mM Tris-HCl, pH 7.5, 150 mM NaCl, 0.5% Triton X-100, 1 mg/ml Aprotinin, 1 mg/ml Leupeptin, 5 mM NaF, 0.5 mM Na_3_VO_4_). The cell lysates were centrifuged at 6000 × g at 4 °C for 15 min. The supernatant was collected as the Triton X-100 soluble fraction, and the pellet was collected as Triton X-100 insoluble fraction. The Triton X-100 soluble fraction was mixed with 6X SDS sample buffer and boiled at 100 °C for 15 min. The Triton X-100 insoluble fraction was cross-linked for 30 min at 37°C with 2 mM disuccinimidyl suberate (DSS) to cross-link the ASC oligomer. After centrifuging at 6000×g for 15 min at 4°C, the pellets were collected and dissolved in 2X SDS sample buffer and subjected to SDS-PAGE.

### Semi-denaturing Detergent Agarose Gel Electrophoresis (SDD-AGE)

Cells were primed with 200 ng/ml LPS for 4 hr, and stimulated by 5 mM ATP or 15 µM nigericin for the indicated duration. Cells were lysed with 1X SDD sample buffer (1X TBE buffer, 10% Glycerol, 2% SDS, 25% Bromophenol blue). NLRP3 oligomers were separated by 1% agarose gel in running buffer (1X TBE and 0.1% SDS) for 1 hr 30 min with 80 V at 4°C. Samples were transferred to the PVDF membrane (Millipore) and detected by using the anti-NLRP3 antibody.

### Two-dimensional polyacrylamide gel electrophoresis (2D-PAGE)

HEK293T cells and MARCH5 knock-downed iBMDMs were plated in 6-well plates. Cells were primed with 200 ng/ml LPS for 4 hr, and followed by 15 µM nigericin for 60 min. Cells were lysed using native lysis buffer (20 mM Tris-HCl pH7.4, 137 mM NaCl, 2 mM EDTA pH8.0, 10% Glycerol, 0.1% Triton X-100, 10 µg/ml Aprotinin, 10 µg/ml Leupeptin, 1 mM PMSF, 0.5 mM NaF, 0.5 mM Na_3_VO_4_) for 15 min on ice. Cell lysates were centrifuged at 20,000×g for 30 min at 4°C, and subjected to 4-12% Blue-Native PAGE. And then, native gels were soaked in 10% SDS for 5 min. For 2D-PAGE, the natively resolved gel was cut well by well and loaded into 4-12% SDS-PAGE gel, followed by conventional western blotting.

### Statistical analysis

All statistical analyses were performed using Prism 6 software (GraphPad). For each result, error bars represent the mean ± SD or mean ± SEM from at least three independent experiments. Statistical significance was measured by a two-tailed unpaired student’s t-test or Mantel-Cox test. P-values are indicated in the figures.

## Data availability

## Acknowledgments

We thanks to Prof. Je-Wook Yu (Yonsei University College of Medicine, Korea) for providing iBMDMs and NLRP3-GFP iBMDMs and, Prof. Ho Chul Kang (Ajou university, Korea) for providing HA-ubiquitin WT and Lys mutants. This work was supported by grant from National Research Foundation of Korea grants funded by the Korean government (MSIP) (NRF-2020R1A2C3011423, NRF-2022R1I1A1A01071281, NRF-2018M3A9H4079660, NRF-2019R1A2C2008283, NRF-2021R1A6A1A03045495).

## Author contributions

**Yeon-Ji Park** : Conceptualization; investigation; formal analysis; methodology; funding acquisition; investigation; investigation; writing-original draft. **Niranjan Dodantenna** : Animal investigation; data curation; formal analysis; investigation; methodology. **Tae-Hwan Kim** : Animal investigation; formal analysis. **Ho-Soo Lee** : Investigation. **Young-Suk Yoo** : Investigation. **Eun-Seo Lee** : Investigation. **Jae-Ho Lee** : Writing – review and editing. **Myung-Hee Kwon** : Data curation; formal analysis. **Ho Chul Kang** : Data curation; formal analysis. **Jong-Soo Lee** : Project administration; supervision; writing – conceptualization; funding acquisition; review and editing. **Hyeseong Cho**: Project administration; supervision; writing – conceptualization; funding acquisition; review and editing.

